# Heat-Triggered Dormancy Release in Low-ROS Pollen Grains Reveals a Conserved Reproductive Reserve

**DOI:** 10.64898/2026.04.30.721981

**Authors:** Asha James, Vinitha Tandle, Nicholas Rutey, Gad Miller

## Abstract

Pollen development and fertilization are considered the most heat-sensitive stages of plant reproduction. While heat stress severely impairs pollen germination and tube growth, the physiological diversity within a single flower’s pollen load suggests that subpopulations may exhibit differential climate resilience. In this study, we tested the hypothesis that this heterogeneity reflects a dormancy-based reserve mechanism that preserves fertilization under heat stress. Using flow cytometry and fluorescence-activated cell sorting in *Arabidopsis thaliana* and *Solanum lycopersicum* (MicroTom), we resolved pollen subpopulations by reactive oxygen species (ROS) status and examined their behavior under increasing heat stress. In both species, ROS-defined metabolic state was tightly associated with pollen size: high-ROS pollen was larger and readily germination-competent, whereas low-ROS pollen was smaller and showed low basal germination, consistent with dormancy. Heat stress preferentially depleted the high-ROS fraction, whereas the low-ROS fraction persisted and, under heat stress, increased metabolic activity and size. By isolating low-ROS and high-ROS pollen, we further show that a brief heat treatment suppresses germination of active high-ROS pollen but promotes germination of dormant low-ROS pollen. These findings provide direct evidence that heat can release dormancy in low-ROS pollen and support a conserved model in which dormant pollen serves as a heat-resilient reproductive reserve.

## Introduction

Dormancy is an adaptive strategy that allows living cells and organisms to preserve viability while postponing growth or development under adverse conditions. Across biological systems, including bacteria, yeasts, animals, and plants, dormant states are marked by reduced metabolism, restricted transcription and translation, and arrested cell cycle and development. At the same time, dormant organisms retain the capacity to rapidly resume activity in response to appropriate environmental or developmental cues. In this way, dormancy serves not only as a survival strategy but also as a means of optimizing the timing of developmental transitions (Marescal and Cheeseman, 2020; Özgüldez and Bulut-Karslioğlu, 2024; Sajeev et al., 2024a).

In pollen, developmental arrest prevails during dispersal. This dormant state is strongly associated with the acquisition of desiccation tolerance, which extends pollen viability during air travel and prevents premature germination before pollen reaches a receptive stigma (Firon et al., 2012; Pacini and Dolferus, 2019).

In many ways, pollen dormancy is similar to seed dormancy. In both cases, dormancy is induced during maturation, preserving the viability of the embryo or gametophyte during desiccation, preventing premature germination, and aligning developmental activation with favorable environmental conditions (Pacini and Dolferus, 2019). Seed dormancy is generally defined as the inhibition of germination in a viable seed under otherwise favorable germination conditions (Chahtane et al., 2016). A similar block appears to exist in pollen, as pollen grains vary in germination capacity, with only a subset readily germinating in vitro under otherwise optimal conditions, while others remain in a dormant, non-germinating state (Boavida and McCormick, 2007; Luria et al., 2019; Zhang et al., 2024).

During seed maturation, the phytohormone abscisic acid (ABA) governs the transition to dormancy and confers desiccation tolerance essential for survival (Finkelstein et al., 2002; Nakashima and Yamaguchi-Shinozaki, 2013; Sajeev et al., 2024b). Whether ABA regulates pollen desiccation and dormancy remains unclear (Sze et al., 2024). Several studies have shown that ABA levels increase during pollen maturation, peaking before anther dehiscence (Chibi et al., 1995; Dai et al., 2018; Li et al., 2021). Transcriptomic analysis also suggests that ABA biosynthesis and signaling contribute to pollen dehydration (Sze et al., 2024).

Interestingly, as in the viviparous seeds of ABA-insensitive mutants (McCarty et al., 1989; Nakashima et al., 2009), several Arabidopsis mutants with defects in specific genes exhibit precocious pollen tube growth within the mother plant’s anthers (Wang et al., 2012; Ju et al., 2016; Zhou et al., 2021). Thus, pollen dormancy is not merely a passive cessation of activity but a regulated physiological state that preserves viability during dispersal and ensures timely activation for fertilization.

Nonetheless, pollen dormancy is clearly distinct from seed dormancy in both timescale and developmental context. Orthodox seeds can remain dry and viable in the soil for years or even centuries, whereas pollen in most angiosperms typically has only hours to a few days to reach a receptive stigma and initiate germination. Structural differences between seeds and pollen may also contribute to the distinct aspects of their dormancy. Seeds are multicellular structures composed of specialized tissues, whereas pollen grains are small bi- or tricellular organisms (Pacini and Dolferus, 2019). Thus, pollen dormancy can be viewed as a short-lived, highly specialized form of developmental arrest, adapted to the male gametophyte’s unique reproductive role. Yet compared with seed dormancy, pollen dormancy remains poorly understood, and the mechanisms that establish, maintain, and release this state are still largely unknown.

The adaptive value of pollen dormancy is likely greatest when environmental conditions challenge pollen survival or disrupt germination timing. This is particularly relevant under heat stress, because the male gametophyte is highly sensitive to elevated temperatures, often resulting in male sterility (Zinn et al., 2010; Santiago and Sharkey, 2019). This sensitivity has made pollen a central focus of recent studies on reproductive thermotolerance.

Because flowers produce far more pollen grains than needed for self-fertilization, reproductive resilience may depend not only on average pollen quality but also on population composition and plasticity. Therefore, understanding pollen thermotolerance requires considering both pollen as a whole and the diversity within pollen populations. Even within a single inbred line, pollen grains can differ markedly in physiological state, including metabolic activity, reactive oxygen species (ROS) production, stress tolerance, and germination potential (Luria et al., 2019; Rutley et al., 2022).

ROS are central regulators of pollen function. They act both as toxic byproducts of metabolism and as signaling molecules that control hydration, germination, and tube elongation (Potocký et al., 2007; Kaya et al., 2014). Using the probe 2’,7’-dichlorodihydrofluorescein diacetate (H_2_DCF-DA), which reports ROS levels in viable cells (as the dye is cleaved by esterases), and flow cytometry, *Luria et al*. (2019) demonstrated that hydrated pollen grains of *Arabidopsis thaliana* and *Solanum lycopersicum* segregate into three reproducible subpopulations based on their ROS-dependent status: no-ROS, low-ROS, and high-ROS. The no-ROS grains, with DCF fluorescence below the detection threshold, are non-viable or metabolically inactive. The low-ROS grains are viable but have low metabolic activity, and the high-ROS grains are fully activated and readily germinate in vitro. This heterogeneity is a constitutive property of mature pollen, reflecting distinct physiological states likely determined by the degree of dormancy established during maturation and desiccation, and by the extent to which this dormancy is maintained after imbibition (Krzyszton et al., 2022; Rutley et al., 2022; Krzyszton et al., 2024).

The level of metabolic activity or dormancy maintenance after imbibition also determines the degree of heat sensitivity. In both Arabidopsis and tomato, moderate heat stress (35°C) sharply depletes the high-ROS pollen fraction, whereas the low-ROS fraction persists or even increases (Luria et al., 2019; Rutley et al., 2021). This differential thermotolerance led us to propose the “pollen in two baskets” strategy, in which dormant, low-ROS pollen serves as a backup for the active, heat-sensitive fraction, thereby preserving fertilization potential after transient stress.

In this study, we tested the hypothesis that dormant pollen serves as a heat-resilient reproductive reserve and extended this conceptual framework. We show that the ROS-defined metabolic state is closely linked to pollen size, with high-ROS pollen larger and low-ROS pollen smaller in both *Arabidopsis thaliana* and *Solanum lycopersicum*. Under heat stress, these subpopulations redistribute dynamically: the active fraction is progressively depleted as temperature rises, whereas the dormant fraction persists and exhibits increased metabolic activity and size. Finally, by isolating dormant and active pollen via FACS, we demonstrate that heat suppresses germination of active pollen while promoting germination of dormant pollen. Together, these findings provide direct evidence that dormant pollen can be reactivated by heat and functions as a stress-resilient reproductive reserve.

## Results

### Pollen metabolic activity is intrinsically coupled to hydration-driven volume expansion in Arabidopsis and tomato

Our previous work on tomato suggested that pollen ROS status is linked to grain size and metabolic state (Rutley et al., 2021). To determine whether this relationship is conserved, Arabidopsis and MicroTom pollen grains stained with the H_2_DCF-DA ROS detector were analyzed by flow cytometry to assess their metabolic status and morphological characteristics at the population level. First, we used the light-scattering properties FSC and SSC as the primary indices of pollen morphology, serving as proxies for grain size and internal complexity, respectively, to characterize the pollen populations.

Initial flow cytometric characterization of Arabidopsis pollen in FSC-A/SSC-A space revealed a seemingly continuous population cloud. However, contour and pseudocolor density visualizations resolved this population into two distinct, partially overlapping clusters, designated P1 and P2 (Fig. 1A–C). This bimodal distribution was even more pronounced in MicroTom tomato, where the two subpopulations exhibited greater centroid separation (Fig. 2A–C). Quantitative analysis of Forward Scatter (FSC-A) indices—a proxy for relative particle size—indicated that P1 consists of smaller grains, whereas P2 represents a larger-diameter population (Figs. 1O and 2O). These physical profiles are morphologically consistent with partially and fully hydrated pollen grains, respectively, suggesting that the populations represent different physiological hydration states at the time of analysis.

**Figure 1.**
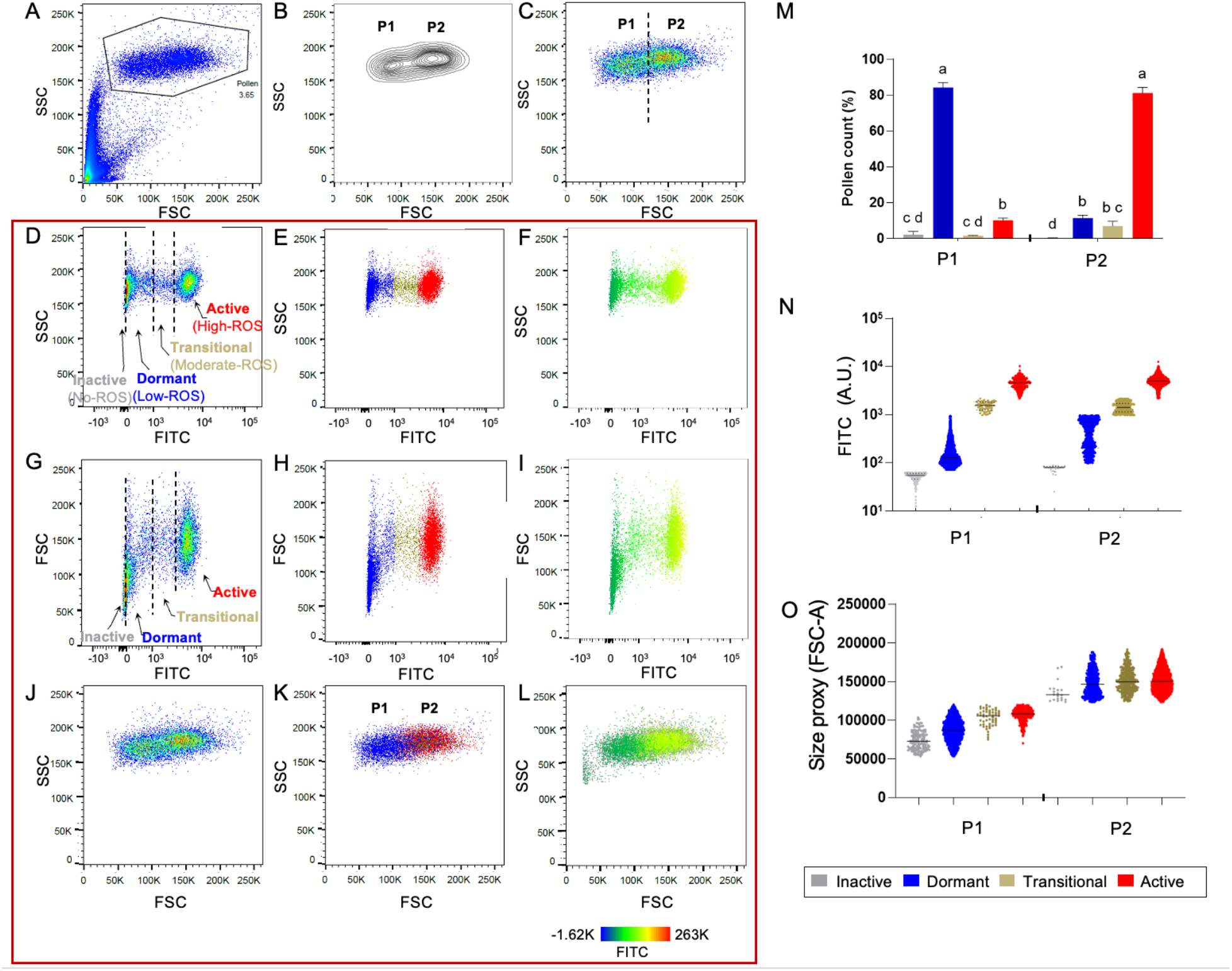
Flow cytometry resolves Arabidopsis pollen into size- and ROS-defined subpopulations. (A) Representative FSC-A versus SSC-A plot of the *Arabidopsis thaliana* pollen population, showing the main gated pollen region. (B, C) Contour and pseudocolor representations of the gated pollen population, resolving two major size classes, P1 and P2, based on forward scatter (FSC-A). (D–F) Representative SSC-A versus FITC fluorescence plots of H_2_DCFDA-stained pollen showing separation into four ROS-defined subpopulations: inactive (no-ROS; gray), dormant (low-ROS; blue), transitional (moderate-ROS; olive), and active (high-ROS; red). (G–I) Representative SSC-A versus FITC showing separation into four ROS-defined subpopulations: inactive (no-ROS; gray), dormant (low-ROS; blue), transitional (moderate-ROS; olive), and active (high-ROS; red). (J–L) Back-gating of inactive, dormant, transitional, and active pollen onto FSC-A versus SSC-A space, showing the position of each subpopulation within the overall pollen population across the P1 and P2 compartments. (M) Relative abundance of the inactive, dormant, transitional, and active pollen fractions within P1 and P2. Bars represent means ± SE of three replicates. Different letters indicate significant differences among groups (two-way ANOVA, Tukey’s test, *P* < 0.05). (N) Quantification of FITC values for the indicated ROS-defined subpopulations and their distribution within the P1 and P2 compartments. (O) Quantification of FSC-A values for the indicated ROS-defined subpopulations, showing that active pollen is enriched in the larger P2 fraction, whereas dormant and inactive pollen are enriched in the smaller P1 fraction. Violin plots represent single-pollen measurements; horizontal lines indicate the median.

**Figure 2.**
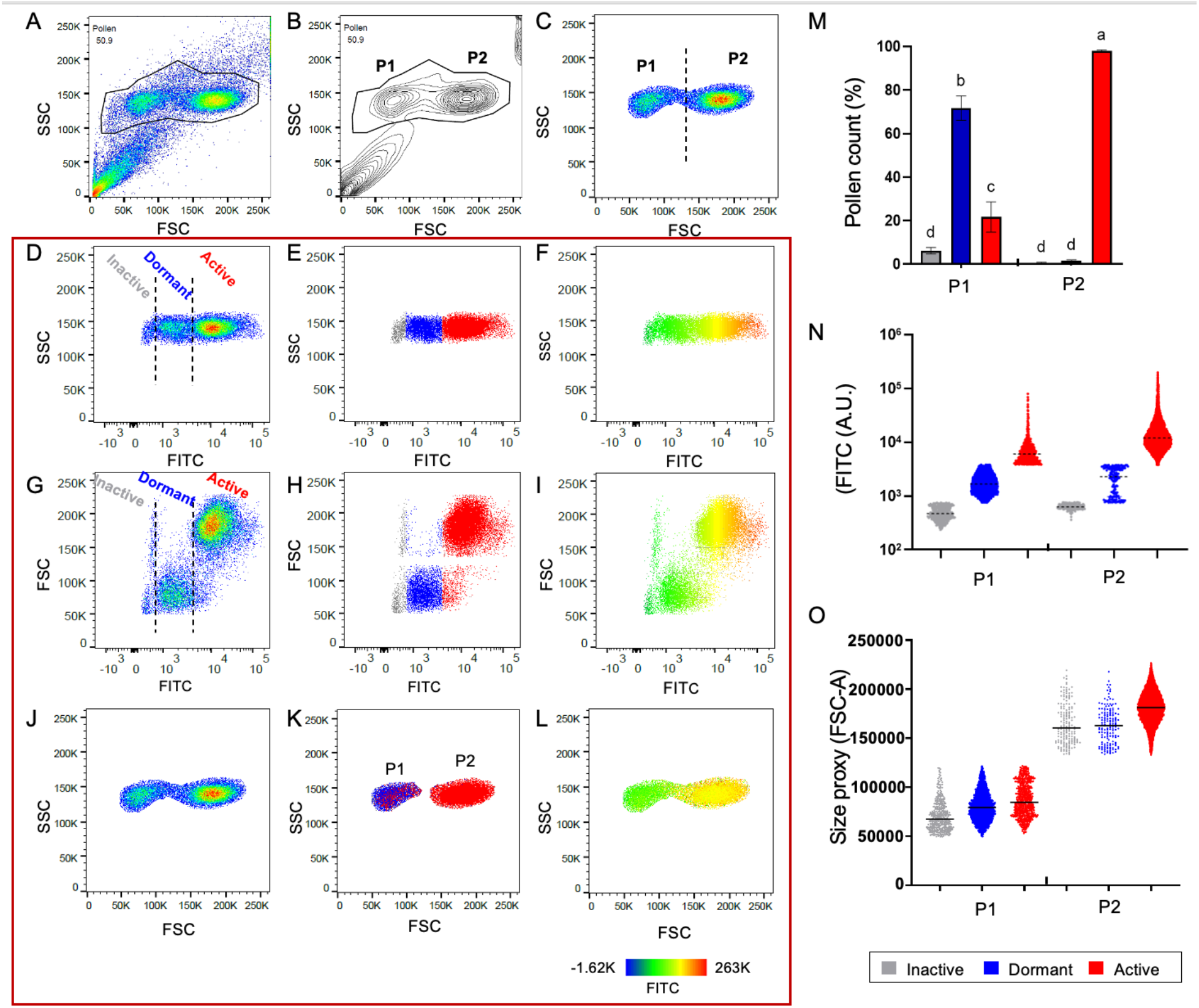
Flow cytometry resolves MicroTom pollen into size- and ROS-defined subpopulations. (A) Representative FSC-A versus SSC-A plot of the *Solanum lycopersicum* cv. MicroTom pollen population, showing the main gated pollen region. (B, C) Contour and pseudocolor representations of the gated pollen population, resolving two major size classes, P1 and P2, based on forward scatter (FSC-A). (D–F) Representative SSC-A versus FITC fluorescence plots of H_2_DCFDA-stained pollen showing separation into three ROS-defined subpopulations: inactive (no-ROS; gray), dormant (low-ROS; blue), and active (high-ROS; red). (G–I) Representative SSC-A versus FITC showing separation into three ROS-defined subpopulations: inactive (no-ROS; gray), dormant (low-ROS; blue), and active (high-ROS; red). (J–L) Back-gating of inactive, dormant, and active pollen onto FSC-A versus SSC-A space, and the position of each subpopulation within the overall pollen population across the P1 and P2 compartments. (M) Relative abundance of the inactive, dormant, and active pollen fractions within P1 and P2. Bars represent means ± SE of three replicates. Different letters indicate significant differences among groups (two-way ANOVA, Tukey’s test, *P* < 0.05). (N) (N) Quantification of FITC values for the indicated ROS-defined subpopulations and their distribution within the P1 and P2 compartments. (O) Quantification of FSC-A values of the indicated ROS-defined subpopulations, showing that active pollen is enriched in the larger P2 fraction, whereas dormant and inactive pollen are enriched in the smaller P1 fraction. Violin plots represent single-pollen measurements; horizontal lines indicate the median.

DCF fluorescence further resolved these populations by their ROS status. In both species, pollen was partitioned into distinct low-ROS and high-ROS fractions, along with a minor unstained (no-ROS) population. In Arabidopsis, the clear separation between the low- and high-ROS peaks revealed a distinct intermediate-ROS (transitional) fraction (Fig. 1D–I). Conversely, the MicroTom distribution was dominated by discrete low- and high-ROS states, with a less prominent transitional population (Fig. 2D–I).

Back-gating these ROS-defined fractions onto the FSC-A/SSC-A space showed a clear correspondence between morphological size class and redox status (Figs. 1K and 2K). In Arabidopsis, the smaller P1 population consisted primarily of low-ROS pollen (84%), with only 11% classified as high-ROS (Fig. 1M). Similarly, in MicroTom, P1 was enriched for the low-ROS fraction (72%), compared with 22% in the high-ROS fraction (Fig. 2M).

The contrast was even more pronounced in the larger P2 populations. In Arabidopsis, P2 was dominated by high-ROS grains (81%), with only 12% in the low-ROS fraction (Fig. 1J–O). In MicroTom, this association was nearly absolute: the P2 cluster consisted of approximately 98% high-ROS pollen, with only 2% in the low-ROS state (Fig. 2J–O).

Collectively, these findings demonstrate a conserved relationship between pollen morphology and physiological state. The enrichment of high-ROS signals in the larger P2 population indicates a transition to a metabolically committed, active state. Conversely, the P1 population maintains a low-ROS redox signature and reduced volume, characteristics consistent with a physiologically quiescent or dormant-like phenotype that may serve as a stress-tolerant reserve.

### Heat stress drives state-dependent physiological shifts in pollen subpopulations

To evaluate subpopulation-specific responses to thermal stress, Arabidopsis and MicroTom pollen were subjected to 30-minute heat treatments across a temperature gradient from moderate to lethal. In Arabidopsis, germination decreased from 95% (at 22°C) to 64% after exposure to 35°C and was completely abolished at 45°C and above (Fig. 3A, E). MicroTom exhibited a lower thermal threshold, with germination declining to 65% at 35°C and failing entirely at ≥40°C. Notably, MicroTom pollen primarily failed by cellular bursting at 40°C, whereas at higher temperatures (45–55°C), grains remained largely intact but ungerminated (Fig. 4A, E). The frequent bursting at 40°C suggests partial metabolic activity without sufficient structural integrity for germination, whereas the stagnant, non-bursting phenotype at ≥45°C indicates a more severe collapse that prevents the turgor increase required for tube emergence.

**Figure 3.**
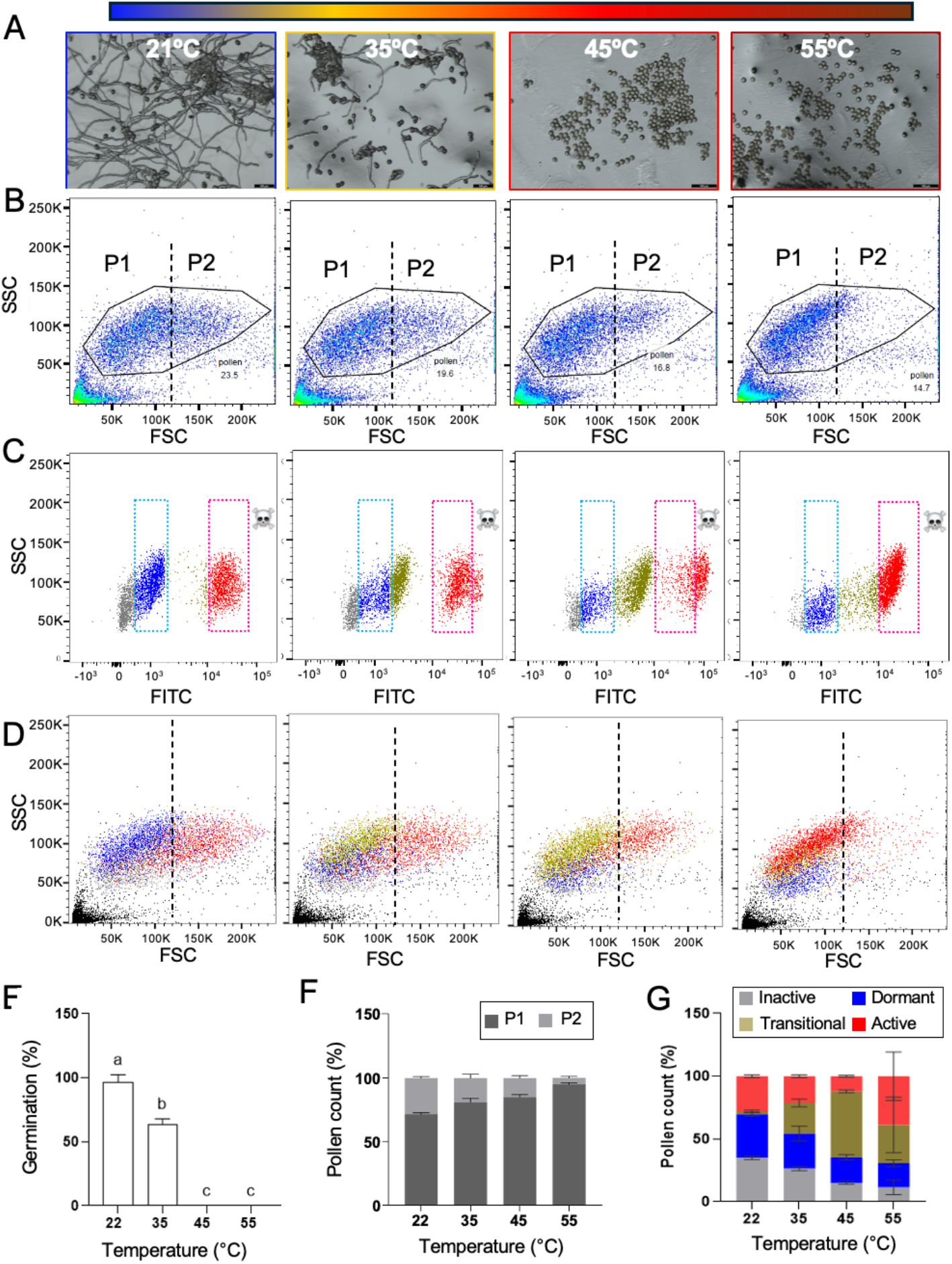
Heat stress progressively reduces Arabidopsis pollen germination and reorganizes size- and ROS-defined pollen subpopulations. (A) Representative bright-field images of *Arabidopsis thaliana* pollen germinated after 30 min incubation at 22, 35, 45, or 55°C, showing progressive reduction in pollen tube formation with increasing temperature. Colored frames correspond to the temperature scale shown above. (B) Representative FSC-A versus SSC-A plots of the total pollen population after heat treatment, showing the gated pollen population and the relative distribution of the small P1 and large P2 fractions. Dashed vertical lines indicate the boundary between P1 and P2. (C) Representative SSC-A versus FITC fluorescence plots of H_2_DCFDA-stained pollen showing the distribution of inactive (gray), dormant (blue), transitional (olive), and active (red) pollen at each temperature. Dashed boxes indicate the primary dormant and active ROS windows. Skull symbols mark regions of excessive ROS levels. (D) Back-gating of ROS-defined fractions onto the FSC-A versus SSC-A space, with the position of each subpopulation within the overall pollen population across the P1 and P2 compartments at increasing temperatures. (E) Quantification of pollen germination after heat treatment, corresponding to (A). (F) Relative abundance of P1 and P2 pollen fractions across temperatures. (G) Relative abundance of inactive, dormant, transitional, and active pollen subpopulations across temperatures. Bars represent means ± SE of three replicates. Different letters indicate significant differences among groups (two-way ANOVA, Tukey’s test, *P* < 0.05).

**Figure 4.**
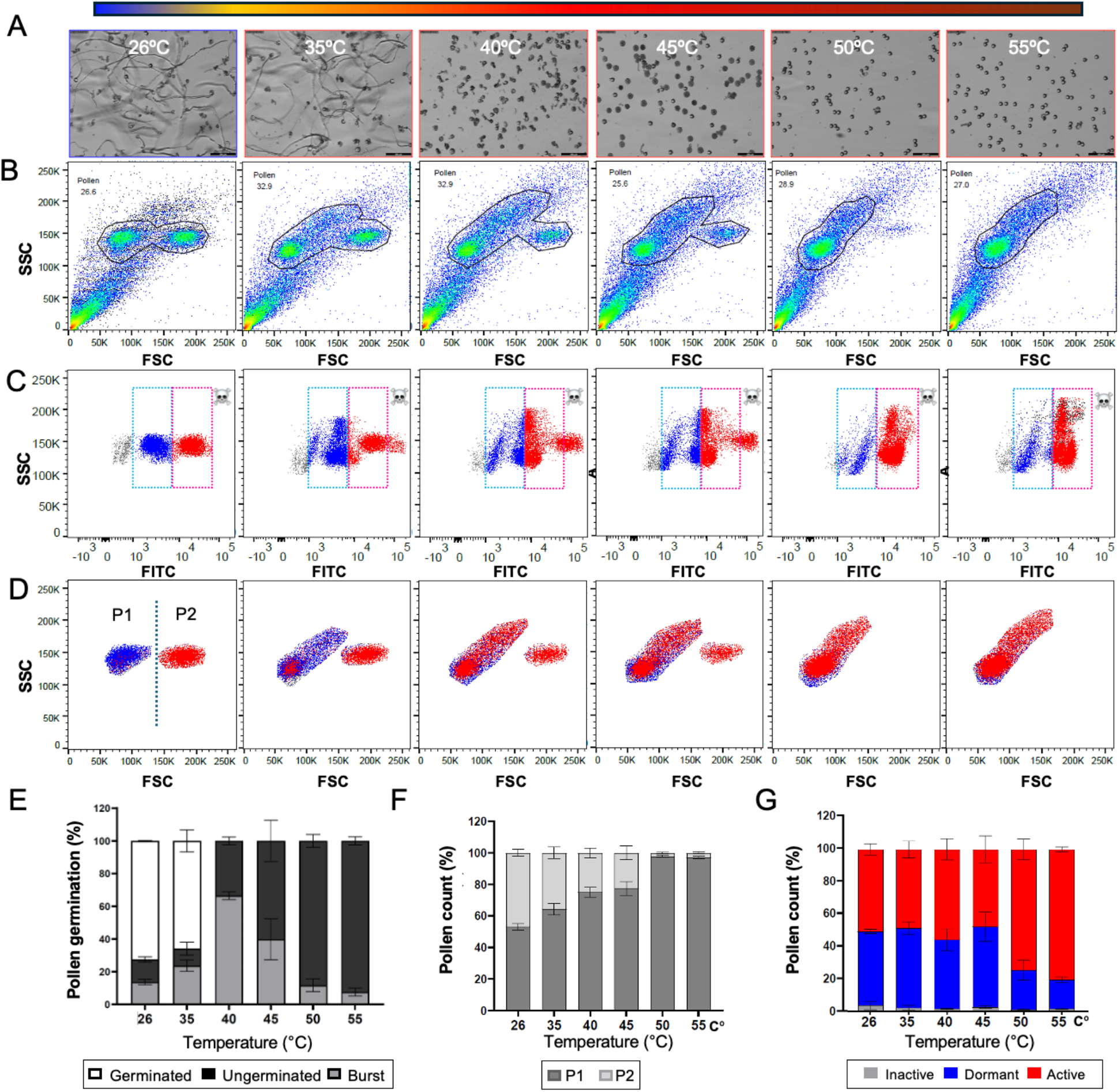
Heat stress progressively eliminates MicroTom pollen germination and reorganizes size- and ROS-defined pollen subpopulations. (A) Representative bright-field images of *Solanum lycopersicum* cv. MicroTom pollen germinated after 30 min incubation at 26, 35, 40, 45, 50, or 55°C, showing normal tube growth under control conditions, increased bursting at moderate heat, and complete loss of germination at higher temperatures. Colored frames correspond to the temperature scale shown above. (B) Representative FSC-A versus SSC-A plots of the total pollen population after heat treatment, showing the gated pollen population and the relative distribution of the small P1 and large P2 fractions. (C) Representative SSC-A versus FITC fluorescence plots of H_2_DCFDA-stained pollen showing inactive (gray), dormant (blue), and active (red) pollen at each temperature. Dashed boxes indicate the principal dormant and active ROS windows. Skull symbols mark regions of excessive ROS levels. (D) Back-gating of ROS-defined fractions onto the FSC-A versus SSC-A space, with the position of each subpopulation within the overall pollen population across the P1 and P2 compartments at increasing temperatures. (E) Quantification of pollen germination outcomes after heat treatment, showing the proportions of germinated, ungerminated, and burst pollen, corresponding to (A). (F) Relative abundance of P1 and P2 pollen fractions across temperatures. (G) Relative abundance of inactive, dormant, and active pollen subpopulations across temperatures. Bars represent means ± SE of three replicates. Different letters indicate significant differences among groups (two-way ANOVA, Tukey’s test, *P* < 0.05).

Flow cytometric profiling revealed a progressive decline in the large P2 population as temperatures increased, suggesting preferential elimination of the larger, ROS-active fraction (Fig. 3B–D, F; Fig. 4B–D, F). Concurrently, the surviving population exhibited a significant rightward shift in DCF fluorescence. In Arabidopsis, this was marked by the emergence of a distinct intermediate-ROS population at 35–45°C, which then transitioned toward the high-ROS gate at 55°C (Fig. 3C, G). In MicroTom, the transition appeared immediate; the surviving grains directly exited the low-ROS window and entered the high-ROS region as the original P2 population vanished (Fig. 4C, G). Analysis of FITC and FSC values further resolved the heat-induced reorganization of pollen subpopulations (Supplemental Figures 1 and 2). Across both species, increasing temperature is accompanied by a progressive redistribution of ROS-defined subpopulations within P1 and P2, along with changes in their relative size (i.e., FSC values). In P1, the inactive fraction decreases gradually, while dormant and active fractions become more prominent, consistent with a stepwise shift from inactive to dormant to active states. A similar succession occurs in P2, where the surviving fractions are progressively reorganized and their relative FSC values shift with heat. This pattern is most evident in MicroTom at 50–55°C, where the surviving pollen is dominated by dormant- and active-like fractions, and the large-pollen P2 compartment appears increasingly replenished by pollen transitioning from P1. Overall, these results demonstrate that heat stress exerts bimodal effects: it selectively compromises the pre-existing active fraction while gradually transitioning the surviving low-ROS population to a higher ROS state. However, because prolonged heat ultimately neutralized germination, this fluorescence shift alone does not distinguish between productive physiological activation and progression toward terminal oxidative toxicity.

### Brief heat stress triggers a reversal of germination competence between active and dormant pollen cohorts

To determine whether the observed shifts in ROS status reflected a productive physiological transition rather than merely cumulative damage, we used FACS to isolate pure cohorts of low-ROS (P1) and high-ROS (P2) pollen. These fractions were subjected to a sublethal thermal pulse (40°C for 7 min in Arabidopsis and 38°C for 8 min in MicroTom) and then assayed for germination competence using a stigma-on-media approach (Fig. 5). Under control conditions (22°C), the two populations exhibited markedly different behaviors. High-ROS grains germinated with relatively high efficiency in Arabidopsis (∼75%) and less so in MicroTom (33%), whereas low-ROS grains remained almost entirely quiescent (0% germination in Arabidopsis; 9% in MicroTom). Strikingly, the sublethal heat pulse induced a complete functional reversal. After heat treatment, the high-ROS population was effectively sterilized, with germination plummeting to <5% in Arabidopsis and 0% in MicroTom. Conversely, the previously dormant low-ROS populations were successfully activated, with germination rates surging to >85% in both species (Fig. 5B).

**Figure 5.**
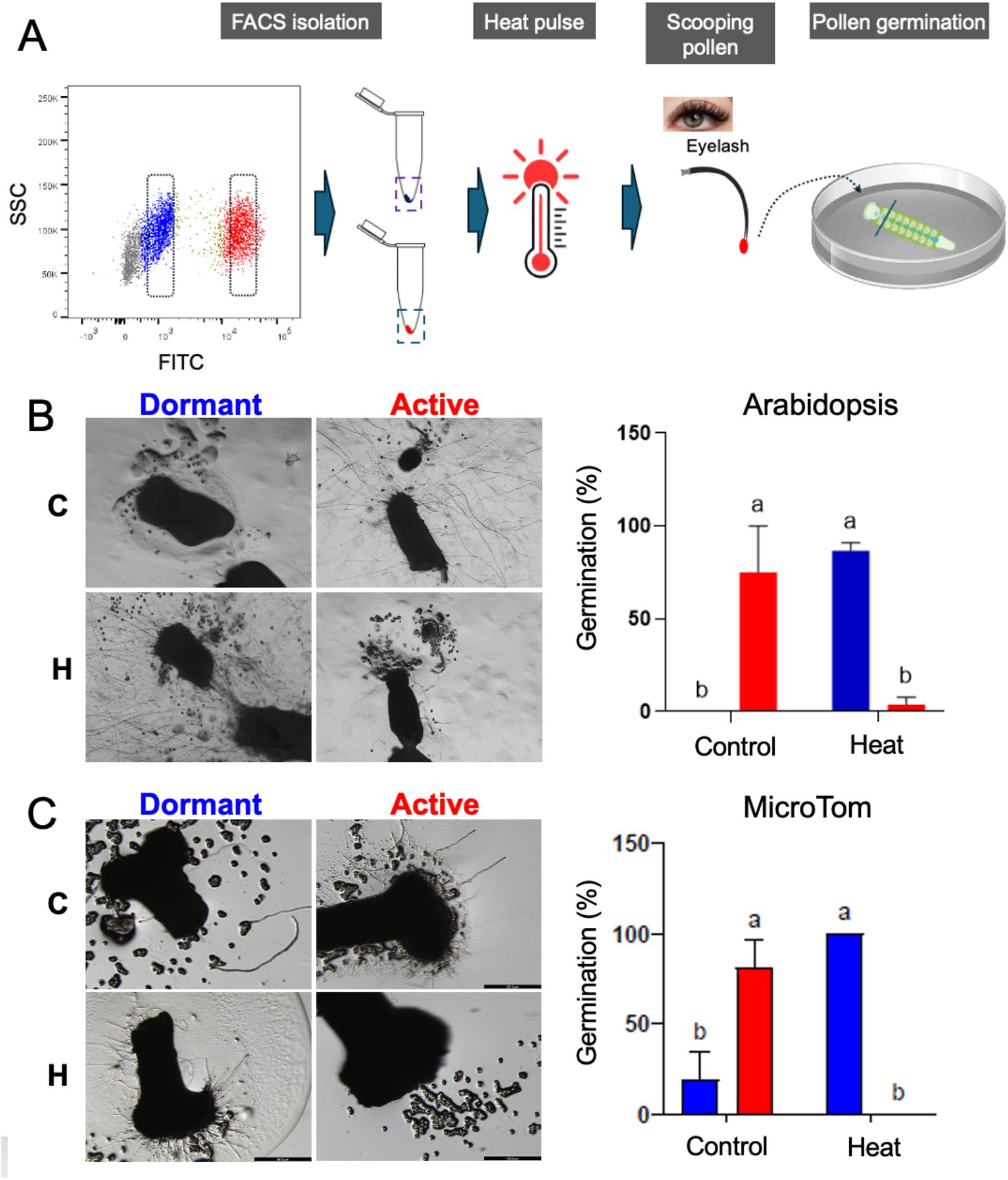
Heat pulse reverses germination competence of dormant and active pollen isolated by fluorescence-activated cell sorting. (A) Schematic representation of the experimental workflow. Dormant (low-ROS) and active (high-ROS) pollen were isolated by fluorescence-activated cell sorting (FACS), exposed to a brief heat pulse, manually transferred using an eyelash tool, and placed on excised pistils positioned on solid germination medium for germination scoring. (B) Representative images and quantification of germination of FACS-isolated dormant and active *Arabidopsis thaliana* pollen under control conditions (C) or after heat treatment (H; 7 min at 40°C). (C) Representative images and quantification of germination of FACS-isolated dormant and active MicroTom pollen under control conditions (C) or after heat treatment (H; 8 min at 38°C). The histograms show the percent germination for dormant (blue) and active (red) pollen. Bars represent means ± SE of three replicates. Different letters indicate significant differences among groups (two-way ANOVA, Tukey’s test, *P* < 0.05).

These data provide functional evidence that pollen thermotolerance is state-dependent. While the metabolically active (high-ROS) fraction is hypersensitive to thermal stress, the dormant (low-ROS) fraction serves as a thermotolerant reserve.

## Discussion

Building on our previous demonstration that pollen populations are physiologically heterogeneous (Luria et al., 2019), this study identifies the low-ROS fraction as a dormant, recruitable subpopulation. Across both Arabidopsis and MicroTom tomato, low-ROS pollen showed minimal germination under control conditions but remained viable and became highly germination-competent after a brief sublethal heat pulse. This reversibility confirms that low-ROS pollen is not inherently defective or damaged; rather, it occupies a restrained physiological state that can be activated by specific environmental cues. These findings assign a clear functional role to this subpopulation, supporting a model in which dormant pollen serves as a resilient reserve during thermal stress.

These results refine our understanding of ROS in pollen biology. Rather than serving merely as a marker of oxidative stress, ROS appears to be a determinant of physiological state: low ROS levels maintain dormancy, a controlled increase accompanies metabolic activation, and excessive accumulation leads to injury. This framework explains why ROS is linked to germination competence under permissive conditions but to toxicity under severe heat stress.

Our observations provide functional evidence for the “pollen in two baskets” model (Rutley et al., 2022), in which population heterogeneity serves as a functional hedge against environmental volatility. Under favorable conditions, the high-ROS fraction is poised for rapid germination. We show that this metabolic readiness is intrinsically coupled to hydration-driven expansion, with the transition from the quiescent P1 state to the active P2 state marked by a significant increase in grain volume. Consequently, Forward Scatter (FSC) serves as a reliable, label-free criterion for distinguishing these cohorts based on intrinsic physical properties. This enables identification of dormant vs. active states independent of external fluorescent markers. This link between physical dimensions and physiological state mirrors a fundamental principle recently established in seed biology. In Arabidopsis, seed size heterogeneity mirrors dormancy depth, with smaller seeds exhibiting higher dormancy and larger seeds possessing greater germination competence (Krzyszton et al., 2024). Our data suggest that size-dependent physiological partitioning is a conserved strategy across plant reproductive units. In this framework, the readiness of the larger, hydrated P2 fraction comes at the cost of extreme heat sensitivity. In contrast, the smaller, low-ROS P1 fraction remains initially restrained but withstands thermal stress more effectively. Once the active fraction is compromised, this thermotolerant reserve can be recruited to re-enter a germination-competent state (Figure 6). Thus, pollen thermotolerance is not merely an average of individual grain survival but an emergent property of the balance between these two distinct morpho-physiological states.

**Figure 6.**
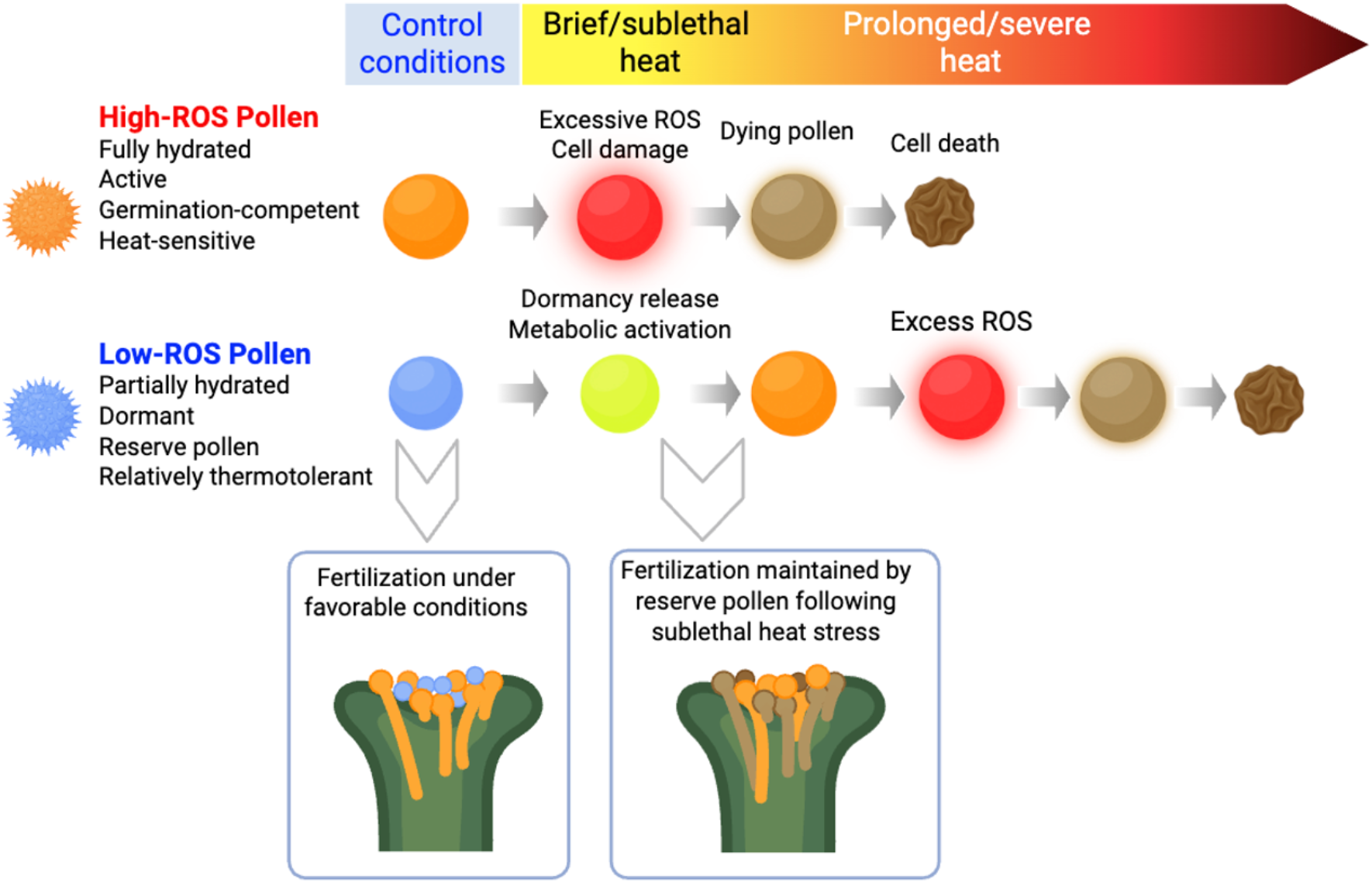
Model of heat-triggered dormancy release and reserve-pollen function under thermal stress. Schematic summary of the relationship among pollen ROS status, hydration state, and fate under increasing heat stress. Under control conditions, active (high-ROS) pollen is fully hydrated, metabolically active, germination-competent, and heat-sensitive, whereas dormant (low-ROS) pollen is partially hydrated, relatively thermotolerant, and serves as a reserve population. A brief heat pulse or sublethal heat dose promotes dormancy release and metabolic activation of dormant pollen, enabling this reserve fraction to contribute to fertilization when the active fraction is compromised. In contrast, active pollen rapidly accumulates excessive ROS under heat stress, undergoes cellular damage, and ultimately dies. Under prolonged or severe heat, the initially activated reserve pollen also accumulates excessive ROS and ultimately loses viability. Insets illustrate the functional outcome of this model: under favorable conditions, fertilization is supported mainly by active pollen, whereas after sublethal heat stress, fertilization can be maintained by reserve pollen.

The conservation of this response pattern in both *Arabidopsis* and *MicroTom*—despite differences in their thermal thresholds—suggests that heat-releasable dormancy is a fundamental feature of pollen biology rather than a species-specific trait.

As in seed dormancy, pollen dormancy maintains viability while restraining germination, metabolic activity remains low, and environmental cues trigger the transition to a growth-competent state (Finch-Savage and Leubner-Metzger, 2006; Sajeev et al., 2024a).

While conceptually analogous to seeds, pollen’s dormant state is uniquely shaped by its gametophytic context and its compressed life cycle (Pacini and Dolferus, 2019).

Unlike orthodox seeds, which may undergo complex cycles of secondary dormancy over multiple seasons (Sajeev et al., 2024a), the narrow window between dispersal and germination in most angiosperms likely requires a more streamlined, one-way activation mechanism. In this context, pollen dormancy primarily prevents precocious germination during transport, preserving metabolic resources until conditions favor successful tube emergence (Wang et al., 2012; Ju et al., 2016; Zhou et al., 2021).

Crucially, we propose that these dynamics define an “oxidative window” for pollen—an adaptation of the redox-gating model established in seed biology (Oracz et al., 2007; Bazin et al., 2011; Oracz and Stawska, 2016). Just as the dormancy-to-germination transition in seeds is governed by crossing a specific redox threshold, our data show that ROS status gates pollen fate: low-ROS grains remain below the activation threshold, high-ROS grains fall within a permissive metabolic window, and those with excessive ROS accumulation exceed the range of physiological viability (Fig. 6).

The physiological basis for the hyperthermosensitivity of active, high-ROS pollen likely lies in pollen’s exceptionally high metabolic rates and mitochondrial densities relative to those of vegetative tissues (Obermeyer et al., 2013). Heat stress can further increase electron leakage from the mitochondrial electron transport chain, triggering ROS overproduction that may rapidly overwhelm the antioxidant capacity of metabolically active grains (Rutley et al., 2021; Suzuki, 2023).

Critically, we demonstrate a discrete thermal window in which heat acts as a physiological activator rather than a proteotoxic stressor, thereby promoting dormancy release and metabolic recruitment. While prolonged heat stress is typically lethal, a transient, sublethal heat pulse can serve as a regulatory cue that stimulates mitochondrial activity and cellular metabolism (Lepock, 2005; Knapp and Huang, 2022). This dual effect aligns with other biological systems in which temperature regulates state transitions (Schulte, 2015; Wendering and Nikoloski, 2023). Correspondingly, a short heat treatment can activate germination of bacterial spores (Wen et al., 2022)and break primary dormancy in various seeds, thereby increasing germination (Piramila, 2012; Luna et al., 2023), a phenomenon we demonstrated to be conserved in pollen. Ultimately, whether a thermal signal results in activation or irreversible injury depends on the thermal dose (temperature and duration) and the pollen grain’s initial physiological state (Fig. 6). Thus, population-level heterogeneity provides a robust mechanism for reproductive buffering, ensuring that a resilient ‘reserve’ of the pollen load remains viable and recruitable amid fluctuating environmental conditions.

Beyond the biological model, this study provides an experimental foundation for a historically underexplored field. By combining ROS-resolved flow cytometry, FACS isolation, and functional germination assays, we have established a tractable system to identify and purify dormant pollen. This establishes a clear path for mechanistic analysis—examining how redox homeostasis, hydration, and heat-shock signaling integrate to regulate the dormant-to-active transition.

Ultimately, reproductive resilience in a warming climate may depend not only on the survival of active pollen but also on the presence of a recruitable dormant reserve. By defining this state across species, we provide a conceptual and methodological framework for dissecting how pollen dormancy contributes to global crop thermotolerance.

## Materials and Methods

### Plant Material and Growth Conditions

***Arabidopsis thaliana*** (accession Col-0) seeds were germinated in soil, and seedlings were transplanted on day 10 into pots containing a soil mixture. Plants were grown under a long-day photoperiod (16 h light/8 h dark) at 21°C. For each experiment, a minimum of 140 plants were used to maximize yield.

***Tomato (Solanum lycopersicum cv. MicroTom)*** plants were grown in controlled-environment chambers at 23°C under a 16 h light/8 h dark photoperiod. Uniform 12-day-old seedlings were transplanted into 15-cm pots and maintained under optimal conditions. Plants were grown under a long-day photoperiod (16 h light/8 h dark) at 24°C.

### Pollen samples collection

Pollen purification, preparation, and analysis were performed as previously described, with some modifications (Rutley and Miller, 2020). Arabidopsis: For flow cytometry and germination assays, freshly opened flowers from 5-6-week-old plants were collected into a 50 mL tube containing pollen germination medium (PGM: 5 mM KCl, 5 mM CaCl_2_, 1 mM MgSO_4_, and 0.01% H_3_BO (pH 7.5)) after vortexing. The extracted pollen grains were filtered through Miracloth (20-25 µm pore size) to remove debris. For FACS experiments, the same extraction protocol was applied, but PGM without sucrose was used.

#### MicroTom

Individual anther cones were collected from freshly opened flowers, with 3 and 6 anther cones per sample for flow cytometry and FACS experiments, respectively, in three replicates per condition. Anther cones were cut transversely, placed in PGM (2 mM HBO_3_, 2 mM Ca(NO_3_)_2_, 2 mM MgSO_4_, 1 mM KNO3, 10% (w/v) sucrose), and gently crushed and vortexed. The released pollen grains were filtered through a 40 µm pore-size nylon mesh to remove debris and collected in 1.5 mL of media per replicate/condition for HS exp and 3 mL of media per replicate for FACS exp.

### In Vitro germination assays for gradual heat stress experiments

Suspended pollen grains were pipetted onto glass slides, which were then covered and placed in a humidified Petri dish. The slides were placed on heat blocks for HS at temperatures ranging from 35 to 55°C for 30 min. After HS, Arabidopsis pollen was incubated overnight at 22°C, and MicroTom pollen for 4 hrs at 26°C. Germination was observed and scored using a Leica stereoscope.

### Dose-response heat stress (HS) experiments

Pollen was hydrated in the PGM for 10 min, then subjected to HS (35-55°C) for 30 min in a pre-heated heat block, and finally stained for ROS analysis. In parallel, control samples were incubated for 30 min at optimal temperatures :22°C for Arabidopsis and 26°C for MicroTom.

### ROS Profiling and Flow Cytometry

Pollen in liquid PGM was stained with H_2_DCFDA (DCF; Cayman Chemical, Ann Arbor, MI, USA) at a final concentration of 2.5 µM for Arabidopsis pollen or 5 µM for MicroTom pollen. H_2_DCFDA stock solutions were prepared in anhydrous DMSO. Staining was performed in the dark to prevent auto-oxidation during incubation of the samples at different temperatures.

Samples were analyzed on a BD LSRFortessa or a BD FACSAria III (BD Biosciences). A minimum of 10,000 events (i.e., pollen grains) was recorded per Arabidopsis sample, and 20,000 per MicroTom sample. Data were processed using FlowJo software (v. 10).

Non-pollen events were excluded using an initial FSC-A/SSC-A gate, and doublets were removed using FSC-A vs. FSC-H gating. The FSC threshold was set to 100V for Arabidopsis and 150V for MicroTom to ensure optimal detection of small P1 grains.

Gating of the high- and low-ROS subpopulations was determined relative to the unstained subpopulation on the FITC-A axis, defining autofluorescence thresholds. FITC-A ranges for the low-ROS subpopulation were gated from 10^2^ to 10^3^, and 10^4^ and above for the high-ROS subpopulation. Gating was slightly adjusted as the subpopulations shifted during sorting. The gated low- and high-ROS subpopulations were then backgated onto the FSC-A vs SSC-A graph to confirm they corresponded to size.

### Fluorescence-Activated Cell Sorting (FACS) experiment

Pollen was extracted as described above using a PGM that does not contain sucrose, then stained with H_2_DCFDA as described above. Two distinct populations (Low-ROS and High-ROS) were resolved and sorted on a BD FACSAria III. The purity of the sorted populations was verified by re-analyzing a small aliquot of each sorted fraction, consistently yielding >95% purity. Pollen isolation - For Arabidopsis and MicroTom pollen, 10,000 and 20,000 grains per fraction, respectively, were sorted into PGM (10% sucrose).

After sorting, samples were concentrated by centrifugation; Arabidopsis pollen at 8,000 rcf for 3 min and MicroTom pollen at 3,000 rcf for 2 min. Pelleted pollen was resuspended to 300 µl, distributed into small Petri dishes, and then subjected to a transient, sublethal HS pulse in a heat block; Arabidopsis: 40°C for 7 min and MicroTom: 38°C for 8 min.

### Post FACS semi-Vivo Germination Assays

After the HS pulse, sorted pollen was carefully transferred to solid PGM using a synthetic eyelash. To stimulate germination under semi-vivo conditions, receptive pistils at anthesis -1 that had been emasculated (Arabidopsis) or freshly opened pistils (MicroTom) were placed on the media. The sorted grains were individually picked with a synthetic eyelash and placed in contact with the stigma (on or near). The plates were then incubated under optimal conditions to recover and maximize germination: Arabidopsis at 22°C for two nights and MicroTom at 26°C overnight in humid chambers.

Three replicates were used for germination scoring and statistical analysis, with a minimum of 30 or 70 grains of Arabidopsis or MicroTom pollen, respectively, per replicate. Experiments were repeated at least three times. Germination was visualized using a Leica stereomicroscope, and a grain was scored as “germinated” if the pollen tube length exceeded the grain diameter.

### Statistical Analysis

All data were analyzed using GraphPad Prism (v. 11). Results are presented as the mean of three technical replicates. Differences in germination percentages were assessed using two-way ANOVA with Tukey’s multiple-comparison test. P-values < 0.05 were considered statistically significant.

## Author Contributions

A.S., V.T., and N.R. performed experiments. A.S. and V.T. analyzed the data. A.S., V.T., N.R., and G.M. designed experiments. G.M. - wrote the original draft of the manuscript. A.S., V.T., and G.M. – prepared the figures. All authors approved the text and provided feedback.

## Supplementary Figure 1

**Supplementary Figure 1. Heat-induced changes in ROS levels** across ROS-defined subpopulations **of Arabidopsis and MocroTom**.

FITC fluorescence intensity of H_2_DCFDA-stained pollen was quantified for the indicated ROS-defined subpopulations in *Arabidopsis thaliana* (top) and (bottom) after exposure to increasing temperatures. In Arabidopsis, FITC values are shown for P1 and P2 inactive, dormant/low-ROS, transitional/medium-ROS, and active/high-ROS fractions at 22, 35, 45, and 55°C. In MicroTom, FITC values are shown for P1 and P2 inactive, dormant, and active fractions at 26, 35, 40, 45, 50, and 55°C. Each violin plot represents the distribution of single-pollen fluorescence values within the indicated gated subpopulation; horizontal black bars indicate medians. The figure highlights the progressive redistribution of pollen among ROS states as temperature increases, including reorganization within both the P1 and P2 fractions. DiYerent letters indicate significant diYerences among subpopulations within each temperature treatment.

**Supplementary Figure 2. Heat-induced changes in pollen size across ROS-defined subpopulations of Arabidopsis and MocroTom**.

Forward scatter area (FSC-A), used as a proxy for relative pollen size/hydration state, was analyzed for the indicated ROS-defined pollen subpopulations in *Arabidopsis thaliana* (top) and MicroTom (bottom) after exposure to increasing temperatures. In Arabidopsis, FSC-A values are shown for P1 and P2 inactive, low-/dormant, medium-/transitional, and high-/active ROS fractions at 22, 35, 45, and 55°C. In MicroTom, FSC-A values are shown for P1 and P2 inactive, dormant, and active fractions at 26, 35, 40, 45, 50, and 55°C. Each violin plot represents the distribution of single-pollen FSC-A values within the indicated gated subpopulation; horizontal black bars indicate medians. Overall, P2 fractions remained larger than the corresponding P1 fractions across temperatures in both species, whereas heat stress progressively altered the relative FSC distribution within and between ROS-defined subpopulations, particularly in the P2 compartment. DiYerent letters indicate significant diYerences among subpopulations within each temperature treatment.

