## Supplemental Figures 1 and 2 for "Heat-Triggered Dormancy Release in Low-ROS Pollen Grains Reveals a Conserved Reproductive Reserve"

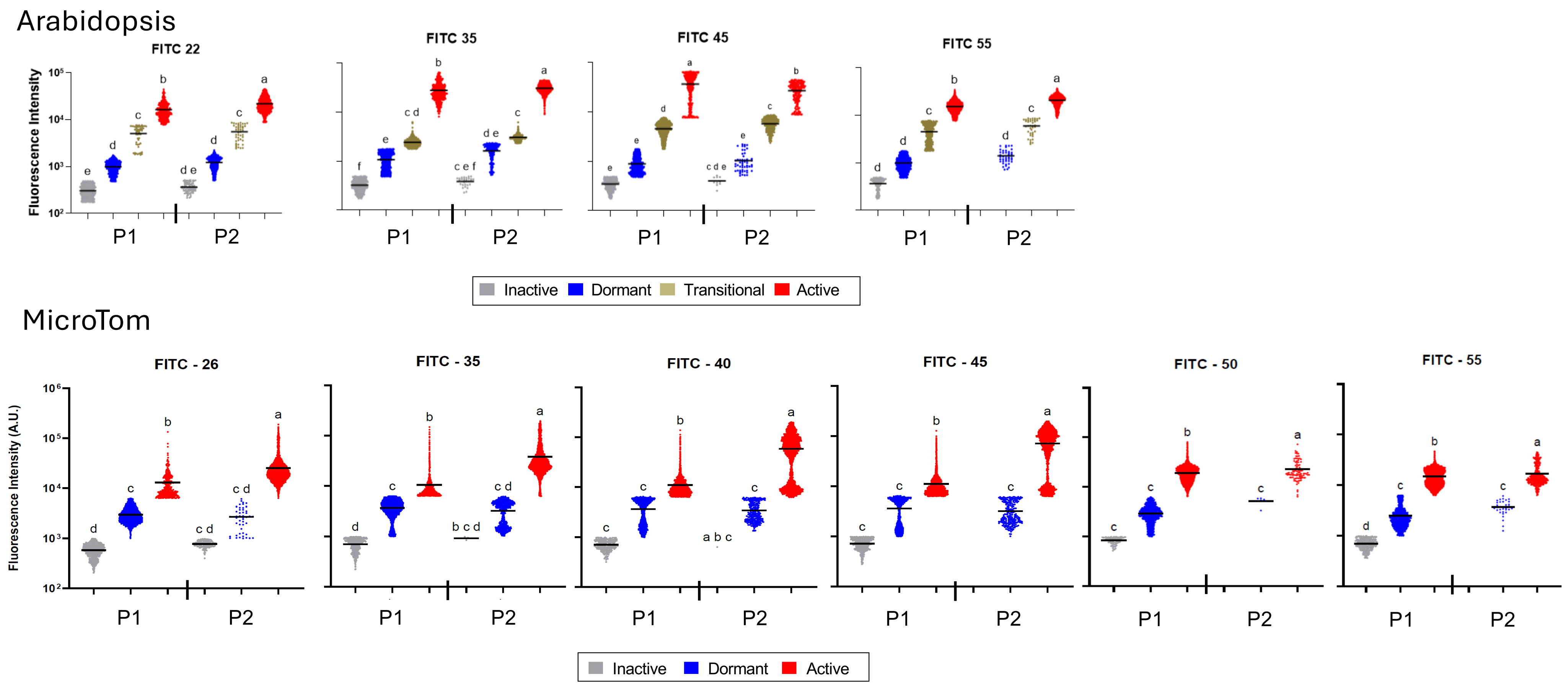


**Supplementary Figure 1. Heat-induced changes in ROS levels** across ROS-defined subpopulations **of Arabidopsis and MocroTom.**
FITC fluorescence intensity of $H_2DCFDA$-stained pollen was quantified for the indicated ROS-defined subpopulations in *Arabidopsis thaliana* (top) and (bottom) after exposure to increasing temperatures. In Arabidopsis, FITC values are shown for P1 and P2 inactive, dormant/low-ROS, transitional/medium-ROS, and active/high-ROS fractions at 22, 35, 45, and 55°C. In MicroTom, FITC values are shown for P1 and P2 inactive, dormant, and active fractions at 26, 35, 40, 45, 50, and 55°C. Each violin plot represents the distribution of single-pollen fluorescence values within the indicated gated subpopulation; horizontal black bars indicate medians. The figure highlights the progressive redistribution of pollen among ROS states as temperature increases, including reorganization within both the P1 and P2 fractions. Different letters indicate significant differences among subpopulations within each temperature treatment.

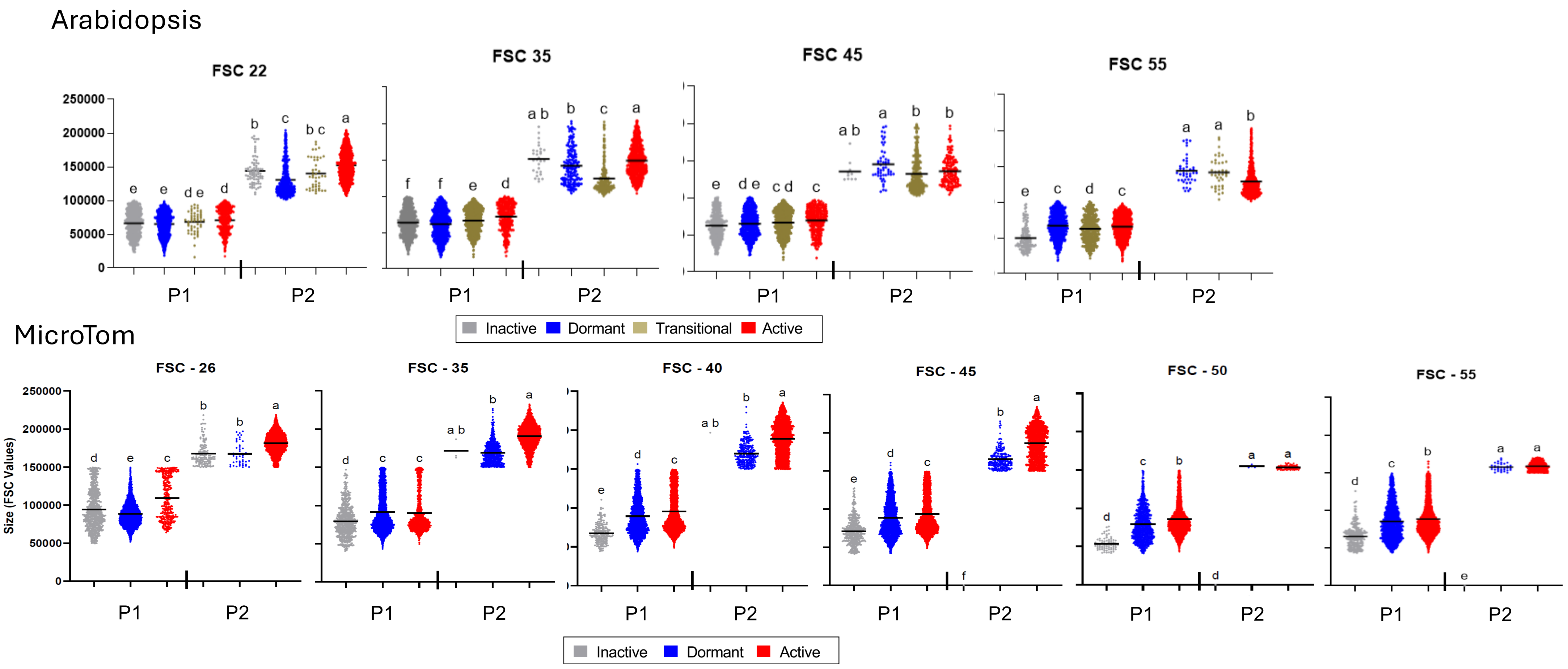


**Supplementary Figure 2. Heat-induced changes in pollen size across ROS-defined subpopulations of Arabidopsis and MocroTom.**
Forward scatter area (FSC-A), used as a proxy for relative pollen size/hydration state, was analyzed for the indicated ROS-defined pollen subpopulations in *Arabidopsis thaliana* (top) and MicroTom (bottom) after exposure to increasing temperatures. In Arabidopsis, FSC-A values are shown for P1 and P2 inactive, low-/dormant, medium-/transitional, and high-/active ROS fractions at 22, 35, 45, and 55°C. In MicroTom, FSC-A values are shown for P1 and P2 inactive, dormant, and active fractions at 26, 35, 40, 45, 50, and 55°C. Each violin plot represents the distribution of single-pollen FSC-A values within the indicated gated subpopulation; horizontal black bars indicate medians. Overall, P2 fractions remained larger than the corresponding P1 fractions across temperatures in both species, whereas heat stress progressively altered the relative FSC distribution within and between ROS-defined subpopulations, particularly in the P2 compartment. Different letters indicate significant differences among subpopulations within each temperature treatment.
